# Polygenic Prediction of Molecular Traits using Large-Scale Meta-analysis Summary Statistics

**DOI:** 10.1101/2022.11.23.517213

**Authors:** Oliver Pain, Zachary Gerring, Eske Derks, Naomi R. Wray, Alexander Gusev, Ammar Al-Chalabi

## Abstract

**Introduction:** Transcriptome-wide association study (TWAS) integrates expression quantitative trait loci (eQTL) data with genome-wide association study (GWAS) results to infer differential expression. TWAS uses multi-variant models trained using individual-level genotype-expression datasets, but methodological development is required for TWAS to utilise larger eQTL summary statistics.

**Methods:** TWAS models predicting gene expression were derived using blood-based eQTL summary statistics from eQTLGen, the Young Finns Study (YFS), and MetaBrain. Summary statistic polygenic scoring methods were used to derive TWAS models, evaluating their predictive utility in GTEx v8. We investigated gene inclusion criteria and omnibus tests for aggregating TWAS associations for a given gene. We performed a schizophrenia TWAS using summary statistic-based TWAS models, comparing results to existing resources and methods.

**Results:** TWAS models derived using eQTL summary statistics performed comparably to models derived using individual-level data. Multi-variant TWAS models significantly improved prediction over single variant models for 8.6% of genes. TWAS models derived using eQTLGen summary statistics significantly improved prediction over models derived using a smaller individual-level dataset. The eQTLGen-based schizophrenia TWAS, using the ACAT omnibus test to aggregate associations for each gene, identified novel significant and colocalised associations compared to summary-based mendelian randomisation (SMR) and SMR-multi.

**Conclusions:** Using multi-variant TWAS models and larger eQTL summary statistic datasets can improve power to detect differential expression associations. We provide TWAS models based on eQTLGen and MetaBrain summary statistics, and software to easily derive and apply summary statistic-based TWAS models based on eQTL and other molecular QTL datasets released in the future.

## Introduction

Genome-wide association studies (GWAS) have identified many replicated genetic associations with complex traits and diseases (1). However, it is challenging to identify the molecular mechanisms underlying these genetic associations. Integration of GWAS summary statistics with functional genomic annotations, such as expression quantitative trait loci (eQTL) data, is a widely used approach for elucidating the molecular mechanisms underlying associated genetic variation (2–4). The integration of eQTL data with GWAS is of particular interest as there is a strong enrichment of eQTLs within genome-wide significant loci, indicating these loci are often mediated by altered gene expression (5).

A popular approach for the integration of eQTL data with GWAS summary statistics is called transcriptome-wide association study (TWAS)(6,7). The TWAS approach aims to infer differential expression associated with the GWAS phenotype. TWAS does this by aggregating the effect of genetic variants associated with the phenotype whilst considering each variant’s effect on a given gene’s expression. This gene-based approach improves power to detect association by reducing the multiple testing burden compared to GWAS, and by aggregating genetic effects across variants in a functionally-informed manner. For TWAS, multi-SNP models predicting each gene’s expression are used, herein referred to as ‘TWAS-models’. Traditionally, these TWAS models are trained using individual-level genotype and gene expression data.

Another popular method that integrates GWAS summary statistics with eQTL data is Summary-based Mendelian Randomisation (SMR) (8). Similar to TWAS, SMR also infers whether a gene’s expression is associated with the GWAS phenotype. In contrast to TWAS, SMR aims to provide evidence for a causal role of a given SNP on a trait mediated through gene expression, focussing on the single largest cis-eQTL effect on gene expression. Currently, a key advantage of SMR over TWAS is that it can use eQTL summary statistics, which are more readily available than the individual-level genotype and expression data traditionally required to derive the multi-SNP models used for TWAS. However, often multiple eQTLs exist for a given gene and considering only the largest eQTL effect will reduce the power to detect associations with gene expression compared to TWAS models allowing multiple eQTL effects. While an extension of SMR incorporating multiple eQTL effects has been developed (SMR-multi)(9),further investigation of methods for integrating eQTL summary statistics with GWAS is required.

A major limiting factor of eQTL databases is the small sample size they are often based upon. Recently, international consortia have pooled eQTL summary statistics from multiple cohorts and performed eQTL meta-analyses, thereby improving the estimation of eQTL effects. Two outstanding recent examples of this are eQTLGen (10) and MetaBrain (11). However, these resources can only be used with methods such as SMR, which do not require individual-level data. Methodology for integrating eQTL summary statistics within a TWAS framework must be developed to better utilize these eQTL summary statistic resources.

There are parallels between the generation of TWAS multi-variant models and the generation of polygenic scores. Similarly, as GWAS transitioned from being carried out in a single sample to the meta-analysis of GWAS summary statistics from multiple cohorts, many polygenic scoring methods have been developed that require only GWAS summary statistics instead of individual-level genotype and phenotype data (12–15). With the establishment of large-scale eQTL meta-analyses, only providing summary statistics, a similar transition from the use of individual-level data to summary statistics is now timely for TWAS. The summary statistic-based *p*-value thresholding and clumping polygenic scoring approach was previously evaluated for generating TWAS models but was found to perform worse than penalised regression models based on individual level data (7). However, polygenic scoring methodology has developed considerably in recent years, warranting the utility of summary statistic-based approaches for generating TWAS models to be revaluated.

In this study, we assess a range of methods for deriving TWAS models predicting gene expression from eQTL summary statistics. We compare the predictive utility of summary statistic-based TWAS models to models derived using traditional methods requiring individual-level data. We additionally compare schizophrenia TWAS results generated using models from either eQTL summary statistics or individual-level data. Furthermore, we explore approaches for defining inclusion criteria for genes in TWAS, selecting the single best model for a given gene, and aggregating TWAS associations across models for a given gene.

## Methods

### Samples

#### eQTLGen

The eQTLGen consortium has performed a meta-analysis of eQTL summary statistics from 37 datasets, totalling 31,368 samples of whole-blood (80.4%) and peripheral blood mononuclear cells (19.6%)(10). The majority of samples are reported to be of European ancestry. Further details of each contributing dataset can be found in the original publication. Full cis-eQTL summary statistics were downloaded from the eQTLGen website (see Data Availability), and cis-eQTL summary statistics excluding GTEx were obtained via private communication with eQTLGen. For each gene, the eQTLGen cis-eQTL summary statistics include all variants within 1Mb of gene boundaries. The eQTL summary statistics only include a signed Z-score to indicate the direction and statistical significance of each eQTL. We converted the Z-score of each variant into a BETA and standard error (SE) to improve compatibility with downstream software. We estimated the BETA as (8)

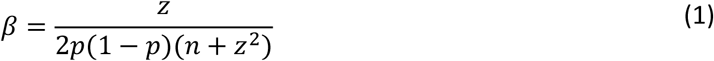

Where *z* is the reported Z-score of the variant in the eQTLGen meta-analysis, *p* is the test allele frequency in the European ancestry subset of the 1000 Genomes (1KG) reference sample (16), and *n* is the sample size for the variant in the eQTLGen meta-analysis. We calculated the SE as

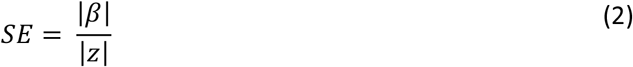

#### YFS

The Young Finns Study collected whole blood from 1,414 individuals across five regions of Finland (17). This dataset has been used several times for TWAS based on TWAS models available on the FUSION website (see URLs). We downloaded the TWAS models from the FUSION website for this study. For each gene, FUSION-released TWAS models include all variants within 0.5Mb of gene boundaries. The FUSION YFS dataset only contains TWAS models for the 4701 genes with a significant SNP-based heritability (*p* < 0.01). We used the same equations as used above for eQTLGen to convert the eQTL Z-score summary statistics within the FUSION top1 models to estimate BETA and SE for each eQTL.

#### GTEx

GTEx (Genotype-Tissue Expression) project release v8 was used as a target sample for evaluating the performance of TWAS models derived using different approaches. GTEx v8 genotype data was downloaded via dbGaP (phs000424.v8.p2) and converted from VCF to PLINK format. The whole blood normalised expression and standard GTEx eQTL analysis covariate data were downloaded from the GTEx portal (see URLs).

### Generating TWAS models from eQTL summary statistics

FUSION is a widely used software for performing TWAS (see URLs)(6). Our study develops an approach concordant with that of FUSION to allow integration with broader TWAS methodology. Consistent with FUSION, SNP-based heritability and TWAS models were generated using HapMap3 (see URLs) variants alone, as these variants are typically well imputed and available in most datasets, improving the portability of the TWAS models to other genotype or GWAS datasets. Variants are not selected based on overlap with specific target datasets. The variants considered for each gene were predetermined by the original study generating eQTL summary statistics or TWAS models.

#### Estimating SNP-based heritability of gene expression

FUSION typically suggests only using TWAS models for genes with a statistically significant SNP-based heritability estimate (*p* < 0.01). This criterion avoids the inclusion of genes for which TWAS models would be unlikely to provide accurate prediction, thereby reducing the multiple testing burden and limiting false positives from null TWAS models. FUSION estimates the SNP-based heritability of gene expression using genome-based restricted maximum likelihood (GREML) and individual-level genotype and gene expression data (6,18).

We evaluated summary statistic-based methods for estimating the SNP-based heritability of gene expression, including linkage disequilibrium score regression (LDSC)(19), and three models implemented within genome-wide complex trait Bayesian analyses (GCTB) software (SBayesR, SBayesR-robust, and SBayesS)(20). SBayesR-robust is the same model as SBayesR but uses an alternative parameterisation procedure which is more robust to misspecification in summary statistics. We included LDSC for comparison with the knowledge that this method is likely to be inaccurate in this context, as only variants within 0.5-1Mb around the gene are considered, whereas LDSC is intended for genome-wide GWAS summary statistics.

We applied these summary statistic methods to eQTL summary statistics derived from the GTEx v8 whole blood FUSION models (European ancestry only) and compared estimates of SNP-based heritability from summary statistics methods to those reported by FUSION estimated using GREML and individual-level GTEx v8 data. The FUSION GTEx v8 models’ SNP-based heritability estimates were available for all genes rather than only genes with significant SNP-based heritability. To avoid excessive computation time, we estimated SNP-based heritability only for genes on chromosome 22 (561 genes).

In addition to SNP-based heritability estimation methods, we also explored a simpler approach for deciding whether genes should be included in the TWAS. We identified genes for inclusion if they had at least one genome-wide significant eQTL present (*p* < 5×10^−8^), or mBAT-combo estimated a significant gene association statistic (FDR < 0.05)(21). The mBAT-combo method combines gene associations from fastBAT (22) and mBAT using the aggregated Cauchy association omnibus test (ACAT-O)(23).

#### Deriving TWAS models predicting gene expression

Models predicting gene expression were derived using several leading summary statistic methods, including SBayesR, SBayesR-robust, LDpred2, lassosum, PRS-CS, DBSLMM, SuSiE and top1 (defined below).

SBayesR, SBayesR-robust (15), LDpred2 (14), lassosum (13), PRS-CS (12) and DBSLMM (24) are reported as leading polygenic scoring methods (25), and can be applied to gene-level eQTL summary statistics. Similar to the methods typically applied within FUSION, these methods use a range of approaches to model genetic effects, allowing for differences in the genetic architecture of genetic effects on gene expression.

LDpred2, lassosum and PRS-CS are often applied using a range of hyperparameters, producing multiple sets of SNP effects. The optimal hyperparameters can then be identified using an external validation sample or estimated using a pseudovalidation method which does not require an external validation sample. Given an external validation sample is often not available, we used the pseudovalidation method for PRS-CS (fully Bayesian model), LDpred2 (auto model), and lassosum (pseudovalidation function) to infer the optimal hyperparameters from the eQTL summary statistics. By default, SBayesR, SBayesR-robust and DBSLMM directly estimate the hyperparameters from the summary statistics, so no additional pseudovalidation step is required. SBayesR and SBayesR-robust were applied using the GCTB-provided precomputed sparse and shrunk LD matrices derived in the European ancestry subset of UK Biobank. LDpred2 was applied using the LDpred2-provided LD matrices also generated in the European ancestry subset of UK Biobank. PRS-CS was applied using the PRS-CS-provided LD matrices generated in the European subset of the 1KG reference. Lassosum and DBSLMM were applied using individual-level data for the European ancestry subset of the 1KG reference.

SuSiE is a method used to perform statistical fine-mapping of GWAS summary statistics to identify variants likely causal for the associated locus (26). SuSiE has also been shown to be useful for generating prediction models, by multiplying each variants’ effects size by the posterior inclusion probability (PIP) estimated by SuSiE (27). We applied SuSiE using default settings, allowing for up to 10 causal signals within each associated locus. SuSiE was applied using LD matrices calculated using PLINK (28) and individual-level data for the European ancestry subset of the 1KG reference.

The model referred to as ‘top1’ simply only considers the variant with the largest absolute Z-score. This model is congruent with the top1 model included by FUSION.

DBSLMM and LDpred2 require an estimate of SNP-based heritability of the gene’s expression. If the SNP-based heritability could not be estimated using the methods above, or the estimate was ≤ 0, we assumed a SNP-based heritability of 0.1 to allow these methods to create models for all genes.

We derived TWAS models using the above methods based on eQTL summary statistics from the full eQTLGen consortium, from the eQTLGen consortium excluding GTEx, and from YFS. We refer to YFS TWAS models from the FUSION website (derived using individual level data) as FUSION-YFS TWAS models and refer the YFS TWAS models derived using eQTL summary statistics as eQTL-YFS TWAS models.

The script used derive to TWAS models from eQTL summary statistics is publicly available (see URLs).

### Evaluating SNP-weights in GTEx v8

We used PLINK (28), implemented by FeaturePred (see URLs), to predict gene expression using all TWAS models in the GTEx v8 target sample. The Pearson correlation between predicted expression and observed whole blood expression was then assessed. The GTEx whole blood normalised expression was adjusted to account for the standard eQTL analysis GTEx covariates, including genetic principal components, surrogate variables, batch, and sex.

We compared the correlation between predicted and observed expression between TWAS models for each gene, determining the statistical significance of differences between models using the

William’s test (also known as the Hotelling-Williams test) (29) as implemented by the ‘psych’ R package’s ‘paired.r’ function, with the correlation between model predictions of each method specified to account for their non-independence. A two-sided test was used when calculating *p*-values.

### Model selection or aggregation methods

Traditionally, TWAS is only performed using the model that explains the most variance in expression of a given gene, as identified by 5-fold cross validation in the training data. However, in the setting of using eQTL summary statistics, formal validation of each model is typically not possible. We explored two alternative approaches to obtain a single indicator of significance for each gene.

#### Pseudovalidation to identify ‘best’ model for each gene

We used the TWAS method to infer which model best predicted expression of a given gene. We adapted the FUSION software (see URLs) to test for an association between all TWAS models and the original eQTL summary statistics. The model with the largest TWAS Z-score for a given gene was assumed to be the most predictive model.

#### Aggregating TWAS results across models

We explored two approaches for aggregating TWAS associations, including FUSION’s omnibus test (FUSION-O) and the aggregated Cauchy association omnibus test (ACAT-O).

FUSION-O is multiple degrees-of-freedom omnibus test, implemented in the FUSION software, which estimates and adjusts for the pairwise correlation between models for a given gene. Default parameters were used, whereby TWAS models are pruned to removed highly correlated models (*R*^2^ > 0.9).

ACAT-O is an aggregated Cauchy association omnibus test, which combines *p*-values without needing to specify the correlation between the models for each gene (23). This method was recently proposed as an omnibus test for TWAS models (30). ACAT-O is robust when combining *p*-values corresponding to different directions of effect, so can be applied to two-sided *p*-values from TWAS.

### Type I error rate

We performed 100 null TWAS to determine the type I error rate (false positive rate) of different strategies for deriving gene associations. Null TWAS results were generated by predicting expression based on the eQTLGen TWAS models into the European ancestry subset of the 1KG reference, and then testing for an association between predicted expression and a random normally distributed phenotype. We measured the type I error rate as the proportion of associations with a *p*-value < 0.05. We measured the type I error rate under several scenarios, selecting gene models using the different criteria, and aggregating TWAS associations across models using different aggregation methods.

### TWAS of schizophrenia

We performed TWAS of schizophrenia using the latest Psychiatric Genomics Consortium schizophrenia GWAS summary statistics (31). We used the summary statistics specific to European ancestry populations (*N*_case_ = 52017, *N*_control_ = 75889) to avoid LD mismatch with eQTL reference datasets. We used applied FUSION-YFS TWAS models, eQTL-YFS TWAS models, and eQTLGen TWAS models (including GTEx v8).

We used and adapted version of FUSION software to perform TWAS testing for association using all TWAS models for each gene (see URLs).

Colocalisation analysis for significant TWAS associations was carried out using the coloc R package (32), as implemented within the FUSION software. This Bayesian approach estimates the posterior probability that associations within a locus for two outcomes are driven by a shared causal variant. It thus enables the distinction between associations driven by pleiotropy (one causal SNP affecting both transcription and MD; posterior probability 4; PP4) and linkage (two causal SNPs in LD affecting transcription and MD separately; posterior probability 3; PP3).

### SMR analysis of schizophrenia

We performed SMR analysis of schizophrenia using the publicly available SMR-formatted eQTLGen data. In contrast to TWAS models, the SMR-formatted eQTLGen data is not restricted to HapMap3 variants. Instead, SMR identifies variants in common between the GWAS and eQTL summary statistics. Therefore, SMR will likely consider variants that are not considered in the TWAS. SMR was run with default parameters, using the European ancestry subset of the 1KG reference sample to estimate LD, restricted to variants with a minor allele frequency > 0.1%.

In addition to SMR, which considers only the most significant eQTL for each gene, we performed SMR-multi, an extension of SMR allowing for multiple eQTL effects on gene expression. SMR-multi was performed simply by adding the ‘--smr-multi’ parameter when running SMR. Default parameters were used, including a *p*-value threshold of 5×10^−8^ to select eQTL for the analysis.

SMR and SMR-multi uses the HEIDI test to determine whether there is effect size heterogeneity between the GWAS and eQTL summary statistics for the given gene. The presence of effect size heterogeneity indicates the association for each trait is being driven by different causal variants. This is conceptually similar to the colocalisation analysis performed within FUSION. Concordant with the original SMR study, we used a HEIDI *p*-value threshold of >0.05 to determine colocalisation of SMR results.

### Extension to MetaBrain eQTL summary statistics

MetaBrain is recently released largescale brain eQTL meta-analysis dataset (11). MetaBrain has released eQTL summary statistics for 5 brain tissues based on European ancestry individuals, including cortex (N = 2,970), basal ganglia (N = 208), spinal cord (N = 108), cerebellum (N = 492), and hippocampus (N = 168). Further details of the eQTL datasets included in MetaBrain can be found in the original publication (11).

After validating our approach for converting eQTL summary statistics into TWAS models using eQTLGen and YFS, we applied the same approach to MetaBrain eQTL summary statistics. Consistent with our analysis of eQTLGen and YFS eQTL summary statistics, for each gene we estimated SNP-based heritability using SBayesR, tested for evidence of eQTL signals using mBAT-combo, derived TWAS models using a range of summary statistic-based polygenic scoring methods, and then performed TWAS of schizophrenia using the MetaBrain TWAS models. We were unable to evaluate the predictive utility of MetaBrain TWAS models in GTEx due to sample overlap.

To determine whether MetaBrain TWAS models can provide novel insights over existing TWAS models based on brain tissue, we compared TWAS results from MetaBrain cortex TWAS models with TWAS results based on PsychENCODE dorsolateral prefrontal cortex (DLPFC) TWAS models (N individuals = 1,321, N genes = 14,751) (33). The PsychENCODE DLPFC sample used to generate TWAS models is smaller than that in the MetaBrain cortex meta-analysis, but the PsychENCODE datasets were combined at the individual-level and the TWAS models were generated using the individual-level based methods implemented within FUSION software. In contrast to the MetaBrain TWAS models, PsychENCODE TWAS models are not restricted to HapMap3 variants, instead considering all high-quality variants after imputation using the haplotype reference panel (34).

## Results

### Predictive utility of TWAS models

TWAS models were generated based on eQTLGen (excluding GTEx) and YFS eQTL summary statistics. TWAS models were then used to predict expression into the independent GTEx v8 sample, and the correlation between the predicted and observed whole blood expression values was calculated. We quantified the number of genes that had a predicted-observed correlation > 0.1, referred to as ‘valid genes’, and we also quantified the number of times each method performed best for a given gene.

When deriving models based on the eQTLGen summary statistics, the method with the highest median correlation and largest number of valid genes was SBayesR-robust (median correlation = 0.045, N valid = 2,449) (Table 1, Figure S1). However, SBayesR-robust was least likely to be the best model for a given gene (6.5%). The discrepancy between the high median correlation but lower relative performance of SBayesR-robust for each gene is due to the method estimating zero SNP-based heritability for 18% of genes, thereby not providing a model for a large proportion of genes. The top1 model was most commonly the best model across genes (20.9%) despite having a relatively low median correlation of 0.035 across all genes. We found each model significantly improved in prediction over all other models for at least 16 genes (Table 1). The top1 model was found to perform significantly worse than the best model for 8.6% of genes in eQTLGen, highlighting the advantage of using models that go beyond the single strongest eQTL for each gene.

**Table 1.** Predicted-observed correlation using eQTLGen TWAS models and GTEx v8 target sample. N valid and % valid indicate the number and percentage of genes that had a predicted-observed correlation > 0.1. N sig. top indicates the number of genes for which a method showed a statistically significant improvement in prediction over all other methods.

| Method | N valid | Median r | % valid | N top | Median r top | % top | N sig. top |
| --- | --- | --- | --- | --- | --- | --- | --- |
| SBayesR-robust | 2449 | 0.045 | 14.60% | 1094 | 0.082 | 6.50% | 22 |
| PRS-CS | 2519 | 0.039 | 15.10% | 1613 | 0.069 | 9.70% | 16 |
| LDpred2 | 2385 | 0.036 | 14.30% | 2156 | 0.061 | 12.90% | 44 |
| DBSLMM | 2116 | 0.036 | 12.70% | 1981 | 0.059 | 11.90% | 22 |
| lassosum | 2385 | 0.035 | 14.30% | 2469 | 0.055 | 14.80% | 20 |
| top1 | 2326 | 0.035 | 13.90% | 3493 | 0.057 | 20.90% | 81 |
| SuSiE | 2003 | 0.033 | 12.00% | 2303 | 0.057 | 13.80% | 24 |
| SBayesR | 1392 | 0.026 | 8.30% | 1603 | 0.049 | 9.60% | 40 |

When using the YFS summary statistics (Table S1, Figure S2), the relative performance of summary statistic methods varied slightly, with PRS-CS models performing best on average. Again, the top1 model performed best for the largest number of genes (25.3%), but the top1 model also performed significantly worse than other models for 12.1% of genes.

We also compared the predictive utility of eQTL-YFS models to FUSION-YFS models, derived using the YFS individual-level data (Table S1). The predictive utility of FUSION-YFS models was typically higher than that of eQTL-YFS models derived using summary statistic-based methods, with FUSION-YFS also typically returning a larger number of valid genes. Despite eQTL-YFS models on average having a lower correlation with observed values, for many genes the eQTL-YFS models performed better than any FUSION-YFS model.

Given the YFS dataset is one of the largest available for TWAS of blood, we tested whether using eQTLGen models significantly improved prediction of expression over FUSION based models for the same genes. Using the FUSION YFS dataset, 4633 genes were imputed into GTEx v8, of which 1,311 had a predicted-observed correlation of 0.1 in GTEx v8. Using the eQTLGen dataset, 16,719 genes were imputed into GTEx v8, of which 3,800 had a predicted-observed correlation of 0.1 in GTEx v8. For genes imputed using both eQTLGen and FUSION YFS datasets (N genes = 4,606), there was a statistically significant increase in the median predicted-observed correlation when using the best eQTLGen model compared to the best YFS model (eQTLGen median *r* = 0.076, YFS median *r* = 0.063, Wilcoxon *p* < 2.2×10^−16^). Figure S3 shows a comparison of predicted-observed correlations for the best model in YFS and eQTLGen. On average, eQTLGen models for genes available in the YFS dataset were more accurate than genes that were not available in the YFS dataset (Figure S4).

### Comparison of eQTLGen TWAS and SMR results for schizophrenia

We compared the results of TWAS and SMR analysis using eQTLGen data. The number of genes tested, the number significant genes, the number of colocalised genes, and the overlap between TWAS and SMR are shown in Table 2.

**Table 2.** Summary of results for eQTLGen based TWAS and SMR analysis of schizophrenia, using different model selection criteria and aggregation methods. N test = Number of genes with non-missing TWAS Z scores in the output; N sig. = Number of significant genes after Bonferroni correction; N sig. overlap with SMR = Number of significant genes also significant in SMR; ObsVal = the model was selected according to the model generated from eQTLGen (excl. GTEx v8) with the highest predicted-observed correlation in the GTEx v8 target sample; Pseudovalidation Z score; PseudoVal = the model was selected with the highest Pseudovalidation Z score; TWAS.P = the model was selected with the smallest TWAS p-value (most significant); top1 = the top1 model was selected; FUSION.O = results for each model was aggregated using the FUSION-O test; ACAT.O = results for each model was aggregated using the ACAT-O test; SNP-h^2^ = only genes with a statistically significant SBayesR SNP-based heritability were retained; mBAT-combo, only genes with either a significant mBAT-combo gene association or genome-wide significant eQTL were retained.

| Approach | N test | N sig. | N sig. coloc. | N sig. overlap with SMR | N sig. coloc. Overlap with SMR |
| --- | --- | --- | --- | --- | --- |
| SMR | 15330 | 217 | 67 | 217 | 67 |
| eQTL (top1) | 18574 | 309 | 105 | 204 | 44 |
| eQTL (ObsVal) | 16562 | 294 | 76 | 147 | 32 |
| eQTL (SNP-$h^2$; ObsVal) | 14225 | 203 | 61 | 102 | 30 |
| eQTL (mBAT-combo; ObsVal) | 14870 | 284 | 76 | 147 | 32 |
| eQTL (PseudoVal) | 19156 | 455 | 90 | 166 | 36 |
| eQTL (SNP-$h^2$; PseudoVal) | 15950 | 342 | 73 | 120 | 34 |
| eQTL (mBAT-combo; PseudoVal) | 16692 | 449 | 90 | 166 | 36 |
| eQTL (ACAT.O) | 19156 | 634 | 115 | 208 | 43 |
| eQTL (SNP-$h^2$; ACAT.O) | 15950 | 443 | 98 | 152 | 42 |
| eQTL (mBAT-combo; ACAT.O) | 16692 | 601 | 116 | 209 | 44 |
| eQTL (FUSION.O) | 19147 | 2179 | 121 | 202 | 39 |
| eQTL (SNP-$h^2$; FUSION.O) | 15856 | 1460 | 101 | 145 | 37 |
| eQTL (mBAT-combo; FUSION.O) | 16684 | 1754 | 122 | 203 | 39 |

As a benchmark for comparison, SMR analysis using eQTLGen eQTL summary statistics identified 217 Bonferroni significant genes, of which 67 passed the HEIDI test indicating colocalisation.

#### Top1 model vs. SMR

We first compared the top1 TWAS models for each gene, as like SMR, the top1 model only considers the strongest eQTL for each gene. Using top1 models, TWAS identified 309 Bonferroni significant genes, of which 204 were significant in the SMR analysis. There was a high Z-score correlation of 0.929 across all genes tested in both TWAS and SMR. We examined the 13 genes that were significant in the SMR analysis but were not significant in the top1 TWAS models. For two of these genes the lead eQTL in the top1 TWAS model was not present in the schizophrenia GWAS, so the TWAS association based on this model was missing. This highlights the advantage of SMR analysing the intersect of the eQTL and GWAS summary statistics, in contrast to TWAS which selects variants for inclusion independent of the GWAS summary statistics. The remaining 11 genes that were significant in SMR but not in TWAS were caused by TWAS using only hapmap3 variants, leading to a difference in the lead eQTL used by SMR and TWAS. The use of different variants between methods typically makes a small difference to the inferred gene expression association due to linkage disequilibrium between variants. However, if there is evidence of effect size heterogeneity between eQTL and GWAS associations within a given locus, using different variants can lead to more substantial differences in the inferred gene expression association. This is demonstrated by the SMR and TWAS (top1 model) Z score correlation increasing to 0.972 among the 441 genes with a HEIDI *p*-value > 0.95 (indicating strong evidence of colocalisation).

#### Model selection and aggregation

We then made comparisons to the TWAS associations for each gene when either selecting the best model using external validation in GTEx v8, selecting the best model using pseudovalidation, or performing TWAS using all models for each gene and then aggregating model specific results.

Simulations showed that the type I error rate was significantly inflated when aggregating associations for each gene using FUSION-O (mean = 0.197, SD = 0.007), but the type I error was well controlled when using ACAT-O (mean = 0.054, SD = 0.003), highlighting that ACAT-O is better calibrated in this context.

TWAS restricted to models externally validated in the GTEx v8 sample resulted in 294 Bonferroni significant genes, of which 76 colocalised with a coloc PP4 > 0.8. Restricting TWAS to genes with external validation in GTEx v8 led to a reduction to the number of genes tested, as 13.5% of the genes in the eQTLGen TWAS models were not present in the GTEx v8 expression data. Evaluation of the pseudovalidation approach to select models in GTEx v8 performed poorly (see Supplementary Information), but TWAS restricted to pseudovalidated models resulted in 455 Bonferroni significant genes, of which 90 colocalised with a PP4 > 0.8. Of the 455 significant genes, 166 were significant in the SMR analysis. Using TWAS with ACAT-O aggregation across models identified 634 Bonferroni significant genes, of which 115 colocalised (Table 2). Of the 634 significant ACAT-O genes, 208 were Bonferroni significant in the SMR analysis. In line with our simulations showing FUSION-O has an inflated type I error rate, when applied to schizophrenia TWAS results FUSION-O identified 2,179 Bonferroni significant genes.

#### Gene inclusion criteria

SNP-based heritability estimation was most accurate when using SBayesR (see Supplementary Information). Restricting the analysis to genes based on statistically significant SNP-based heritability substantially reduced the number of Bonferroni significant genes identified, the number of colocalised genes, and the overlap with significant genes in the SMR analysis. Restricting to genes with a significant mBAT-combo signal or a genome-wide significant eQTL led to 33 fewer Bonferroni significant genes, but the number of significant and colocalised genes increased by one gene. This indicates restricting the analysis to genes with significant evidence of eQTL effects present may reduce the number of associations driven by linkage (different causal variants) and thereby reduces the multiple testing burden to detect colocalised associations at statistical significance.

#### Comparison to SMR-multi

We additionally compared TWAS results to those of SMR-multi, a multi-variant extension to SMR. SMR-multi identified 508 Bonferroni significant associations, of which 108 passed the HEIDI test indicating colocalisation. This result highlights SMR-multi provides a gain in power compared to SMR, and many of the novel associations pass the HEIDI test and are therefore consistent with a causal model.

Of the 508 Bonferroni significant associations, 186 were significant in the SMR analysis, 230 were significant across pseudovalidated TWAS models, and 315 were significant in the ACAT-O aggregated TWAS.

Ultimately, the mBAT-combo-restricted and ACAT-O aggregated TWAS of schizophrenia identified 25 genes as significant and colocalised that were not significant in either SMR or SMR-multi analyses (Table 3).

**Table 3.** Significant and colocalised eQTLGen schizophrenia TWAS associations that were not significant in SMR or SMR-multi. Showing genes with significant mBAT-combo association or genome-wide significant eQTL, that were Bonferroni significant after aggregation across TWAS model using ACAT-O, and colocalised using a coloc PP4 threshold of 0.8. Mean TWAS Z indicates the mean TWAS Z-score across models for each gene to indicate the direction of the association.

| Location | Name | Mean TWAS Z | ACAT-O P (Bonferroni) | Coloc PP4 | SMR P (Bonferroni) | SMR-multi P (Bonferroni) | HEIDI P |
| --- | --- | --- | --- | --- | --- | --- | --- |
| 1:6684926-6695646 | THAP3 | -5.17 | 7.03E-06 | 0.981 | 0.391 | 0.108 | 0.073 |
| 1:8484705-8494898 | RP5-1115A15.1 | -4.10 | 5.08E-04 | 0.968 | 0.131 | 1.000 | 0.001 |
| 1:36179476-36185073 | C1orf216 | 4.43 | 1.09E-03 | 0.896 | 0.139 | 0.061 | 0.778 |
| 1:150237804-150241980 | APH1A | -4.02 | 6.07E-04 | 0.938 | 1.000 | 0.825 | 0.010 |
| 1:160185505-160254920 | DCAF8 | 4.70 | 3.01E-04 | 0.945 | 0.689 | 1.000 | 0.857 |
| 2:71503691-71662199 | ZNF638 | 3.85 | 1.24E-06 | 0.847 | 1.000 | 1.000 | 0.015 |
| 3:9391373-9440263 | SETD5-AS1 | 4.47 | 1.56E-05 | 0.868 | 1.000 | 1.000 | 0.423 |
| 3:63850233-63989138 | ATXN7 | -5.06 | 1.11E-06 | 0.982 | 0.394 | 0.394 | 0.272 |
| 4:156263810-156298122 | MAP9 | 4.12 | 1.12E-03 | 0.978 | 0.623 | 0.222 | 0.444 |
| 6:111804714-111919505 | TRAF3IP2-AS1 | -4.12 | 8.10E-04 | 0.988 | 1.000 | 1.000 | 0.185 |
| 7:100277130-100287071 | GIGYF1 | 4.42 | 3.72E-05 | 0.974 | 0.085 | 1.000 | 0.470 |
| 8:26240414-26363152 | BNIP3L | 4.37 | 1.37E-06 | 0.998 | 1.000 | 1.000 | 0.363 |
| 9:131932713-131933219 | RP11-247A12.7 | 4.58 | 4.74E-05 | 0.861 | 0.089 | 0.358 | 0.604 |
| 10:64571756-64679660 | EGR2 | -3.18 | 6.04E-04 | 0.993 | 1.000 | 1.000 | 0.341 |
| 11:57520715-57587018 | CTNND1 | 3.36 | 7.13E-04 | 0.881 | 0.336 | 1.000 | 0.005 |
| 11:65479467-65487075 | KAT5 | 4.34 | 1.49E-03 | 0.804 | 1.000 | 1.000 | 0.369 |
| 16:4560676-4588829 | CDIP1 | 4.74 | 4.97E-04 | 0.829 | 1.000 | 1.000 | 0.275 |
| 17:5185558-5289129 | RABEP1 | -2.51 | 1.26E-06 | 0.946 | 0.068 | 0.346 | 0.571 |
| 17:12692856-12894960 | ARHGAP44 | -4.36 | 5.32E-04 | 0.891 | 1.000 | 1.000 | 0.018 |
| 17:17713713-17740325 | SREBF1 | -3.40 | 6.96E-04 | 0.867 | 0.180 | 0.486 | 0.150 |
| 17:17942606-17971718 | GID4 | -4.23 | 4.13E-04 | 0.884 | 0.210 | 0.502 | 0.033 |
| 17:43471275-43511787 | ARHGAP27 | 2.48 | 2.32E-05 | 0.992 | 1.000 | 1.000 | 4.47E-06 |
| 17:53246184-53250993 | CTC-462L7.1 | 3.65 | 3.88E-06 | 0.86 | 0.884 | 1.000 | 0.899 |
| 19:50138549-50143458 | RRAS | -5.57 | 1.17E-08 | 0.996 | 0.148 | 0.089 | 0.658 |
| 22:25844072-25916821 | CRYBB2P1 | -4.96 | 2.81E-11 | 0.932 | 1.000 | 1.000 | 0.052 |

### Comparison of individual-level and summary statistic-based YFS TWAS

We compared TWAS results when using models derived using individual-level YFS data (FUSION-YFS models) and models derived using YFS eQTL summary statistics (eQTL-YFS models). The number of genes tested, the number of significant genes, the number of colocalised genes, and the overlap between FUSION and eQTL based TWAS are shown in Table S2.

Across all 4,685 genes, TWAS using FUSION-YFS models, selecting the best model per gene based on cross-validation *r*^2^ (CV.R2, standard TWAS approach), identified 93 Bonferroni significant genes.

TWAS using eQTL-YFS models, selecting the best model using pseudovalidation (PseudoVal), resulted in 75 Bonferroni significant genes, 58 of which were also Bonferroni significant in the FUSION-YFS TWAS. The TWAS Z-score correlation between FUSION-YFS (CV.R2) and eQTL-YFS (PseudoVal) models was 0.851.

Consistent with eQTLGen results, ACAT-O improved power to detect associations, identifying 162 Bonferroni significant genes, 87 of which were Bonferroni significant in the FUSION TWAS. We additionally show that applying ACAT-O to FUSION-YFS TWAS results also increased the number of identified associations, identifying 130 Bonferroni significant associations, indicating this approach improves power to detect associations even when the best TWAS model for each gene has been identified via 5-fold cross-validation.

Similar to eQTLGen results, restricting the analysis to genes with significant SNP-based heritability substantially reduced the number of significant genes identified by TWAS (Table S2), but restricting the analysis to genes with significant mBAT-combo associations or with a genome-wide significant eQTL retained all significant associations.

### MetaBrain TWAS and comparison to PsychENCODE

Based on our previous results using eQTLGen and YFS, for the MetaBrain TWAS, we only included genes with an FDR significant mBAT-combo association or genome-wide significant eQTL, and aggregated TWAS associations across models using ACAT-O. The number of genes retained for each MetaBrain tissue varied depending on the sample size available (Table S4).

We compared the MetaBrain cortex and PsychENCODE TWAS results. TWAS using MetaBrain cortex eQTL summary statistics identified 374 significant genes, of which 59 showed strong evidence of colocalisation. TWAS using PsychENCODE models based on individual-level data, selecting the best model for each gene based on the out-of-sample variance explained, identified 211 significant genes, of which 78 showed strong evidence of colocalisation. Across TWAS using MetaBrain cortex and PsychENCODE DLPFC, 61 genes were identified as statistically significant using both, and 11 showed strong evidence of colocalisation using both. The 374 significant genes in the MetaBrain cortex TWAS were separated into 114 independent loci (0.5Mb window), of which 46 contained a significant gene in the PsychENCODE TWAS, indicating TWAS using MetaBrain identifies associations within novel loci.

These results indicate MetaBrain cortex TWAS and PsychENCODE TWAS often identify different associations, with MetaBrain cortex identifying more significant associations, but fewer colocalised associations.

## Discussion

This study derives and evaluates a TWAS-based framework for integrating eQTL summary statistics with GWAS summary statistics to infer differential gene expression associated with the GWAS phenotype. We use eQTL summary statistics from eQTLGen and YFS datasets to derive TWAS models, evaluating the predictive utility of these models within the independent GTEx v8 target sample. We use the eQTL summary statistic-based TWAS models to perform TWAS of schizophrenia, comparing results to those identified using existing methods and resources.

### Predictive utility of summary statistic-based TWAS models

We show that summary statistic methodology can be applied to eQTL summary statistics to derive multi-variant TWAS models predicting gene expression in external datasets. We demonstrate that multi-variant TWAS models based on eQTL summary statistics often significantly improve prediction of gene expression over the top1 model using only the single largest eQTL, although the best model varies across genes. Furthermore, TWAS models derived using individual-level data provided only a small improvement over eQTL summary statistic-based TWAS models, and as a result, TWAS models using the larger eQTLGen summary statistics provided significantly improved prediction over models based on the YFS individual-level data. This was demonstrated by showing the number of ‘valid’ genes (with predicted-observed correlation > 0.1) increased when using eQTLGen data compared to YFS data, and by showing the predicted-observed correlation for genes increased when using eQTLGen data over YFS data. These findings support the use of multi-variant TWAS models based on larger eQTL summary statistics when corresponding individual-level data is not available.

### TWAS model selection and aggregation

TWAS models can be integrated with GWAS summary statistics to perform TWAS or used to predict gene expression levels in individual-level data. Traditionally, in both these situations, the TWAS model with the highest predictive utility is selected for each gene, determined using cross-validation in the training data. Given we typically do not have a validation dataset to evaluate eQTL summary statistic-based TWAS models, we explored two alternative approaches to either select or aggregate TWAS models.

We evaluated the TWAS method as a form of pseudovalidation, to select the model best predicting gene expression. However, using the same eQTL summary statistics to derive and test the TWAS models appeared to lead to overfitting, with larges biases towards more complex models. Further methodological development is required to infer the optimal model for a given gene using summary statistics alone. In the meantime, we recommend using the best model identified using formal validation, as we have done by evaluating the eQTLGen TWAS models in the independent GTEx v8 target sample.

When performing TWAS, it is possible to aggregate evidence of association across TWAS models for a given gene. We explored two approaches for this, including ACAT-O and FUSION-O. Simulations highlighted the type I error rate was well controlled when using ACAT-O, but not FUSION-O. Using ACAT-O to aggregate associations for each gene identified more significant associations than using either formal validation or pseudovalidation to select the single ‘best’ model, indicating that ACAT-O model aggregation can improve statistical power to detect significant associations. Therefore, when multiple models for each gene can be considered, such as TWAS, model aggregation may be preferable to selecting the single best model.

### TWAS of schizophrenia and comparison to existing resources

We performed a TWAS of schizophrenia using TWAS models derived from eQTLGen summary statistics. Comparison to SMR and SMR-multi showed that using the TWAS approach to integrate eQTL summary statistics with GWAS can identify novel associations over SMR and SMR-multi.

However, SMR and SMR-multi also identified associations that were not significant in the TWAS, indicating that these different approaches can each offer novel insights. Our TWAS, restricted to genes with strong eQTL effects and aggregating using ACAT-O across TWAS models, identified 25 significant and colocalised genes that were not identified using either SMR or SMR-multi. This illustrates that TWAS can provide novel insights over existing methods that can be used to further our understanding of disease aetiology.

We further extend our summary statistic-based approach to MetaBrain eQTL summary statistics, comparing schizophrenia TWAS results to those generated using the smaller but individual-level PsychENCODE dataset. Compared to PsychENCODE, the MetaBrain TWAS identified a larger number of significant associations but fewer associations colocalised. This suggests that MetaBrain TWAS increases power to detect associations over existing resources, but the signals often do not colocalise. One possible explanation for the reduced likelihood of colocalisation is that MetaBrain is a meta-analysis of multiple datasets which often leads to different individuals being considered across variants, which may increase the likelihood of effect size heterogeneity when compared to the GWAS summary statistics. Additionally, many associations were unique to either MetaBrain or PsychENCODE TWAS, indicating TWAS models from both sources can offer novel insights.

### Gene inclusion criteria

Often in there is insufficient data to identify robust eQTL associations for a given gene. Including TWAS models for genes with weak eQTL associations are unlikely to be useful for prediction in external individual-level data or for TWAS analysis, which may introduce noise into downstream analyses and unduly increase the multiple testing burden. Therefore, we explored two approaches for identifying genes with a sufficient eQTL signal present, including statistically significant SNP-based heritability of gene expression, and identifying genes with either a significant mBAT-combo association or genome-wide significant eQTL present. We found that restricting TWAS to genes with statistically significant SNP-based heritability substantially reduced the number of significant and colocalised TWAS associations, indicating this approach reduced the power of TWAS. In contrast, restricting the TWAS to genes with either a significant mBAT-combo association or genome-wide significant eQTL present retained most significant TWAS associations, and slightly increased the number of colocalised genes, highlighting that this approach is a useful criterion when selecting TWAS models for downstream analysis.

### Comparison with OTTERS

A recent preprint describes a similar approach to ours for the integration of eQTL summary statistics in a TWAS framework, called OTTERS (30). Our findings are congruent, with summary statistic methods applied to eQTLGen providing novel opportunities and associations. Notable advances made by our study are as follows. First, in addition to lassosum and PRS-CS models for gene expression we include DBSLMM, LDpred2, SBayesR, SBayesR-robust, and SuSiE. In contrast OTTERS includes two alternative methods including the *p*-value thresholding and clumping approach, and SDPR. Second, we explore criteria for including genes in TWAS, including SNP-based heritability and the mBAT-combo gene association test, with the latter avoiding the inclusion of many genes with invalid prediction. OTTERS generates models for all genes, potentially including models with no predictive utility. Third, our approach for deriving TWAS models is very fast, taking 1-3 minutes per gene, in contrast to the ∼20 minutes reported in the OTTERS preprint. Fifth, our study provides additional comparisons with existing methods and resources, providing a comparison of methods to existing TWAS models from the YFS sample, and providing a comparison to eQTLGen results derived using SMR and SMR-multi.

### Limitations

There are limitations to the application of eQTL summary statistics to the TWAS framework. Most summary statistic methods require the use of an LD reference, matched to the ancestry of the population used to generate the summary statistics. Misspecification of LD reduces the accuracy of summary statistic-based methods, leading to reduced accuracy of prediction models. Furthermore, summary statistic-based methods, such as colocalisation are sensitive to misspecification that can occur when using summary statistics from largescale meta-analyses. Integration of methods that attempt to resolve these LD mismatch and misspecification issues may further improve the value of eQTL summary statistic datasets (35,36). A further limitation of using eQTL summary statistics is the absence of a validation sample to identify the optimal hyperparameters or model for a given gene, and the current inaccuracy of our proposed pseudovalidation approach. Although we provide formal validation results for the eQTLGen TWAS models derived in this study, further investigation of pseudovalidation methods should be explored that diminish the likelihood of overfitting for future eQTL summary statistic datasets.

### Implications

We have developed our approach for the integration of eQTL summary statistics in line with the existing and popular FUSION software for performing TWAS. This allows others to easily integrate TWAS models from novel eQTL summary statistics resources. We have made the TWAS models derived from eQTLGen and MetaBrain publicly available for others to use (see Data Availability). The scripts used to define gene inclusion criteria and compute prediction weights is also highly efficient, taking 1-3 minutes per gene, and can be run in parallel for each gene, allowing other researchers to easily compute TWAS models from eQTL summary statistics as they are released.

GWAS methodology transitioned from the use of individual-level data to summary statistics to overcome data sharing restrictions and increase the sample size and statistical power. Our study facilitates a similar transition, specifically for methodology applied to eQTL data and other molecular QTL data (e.g., methylation, protein). The application of summary statistic-based methods to eQTL summary statistics provides novel and enhanced opportunities for the utilisation of eQTL summary statistics to understand and predict complex phenotypes.

## Data availability

- TWAS models from eQTLGen summary statistics and corresponding validation results in GTEx v8: https://doi.org/10.5281/zenodo.7068381
- TWAS models from MetaBrain summary statistics: https://doi.org/10.5281/zenodo.7121234
- YFS TWAS models: http://gusevlab.org/projects/fusion/
- eQTLGen cis-eQTL summary statistics: https://www.eqtlgen.org/cis-eqtls.html
- Schizophrenia GWAS summary statistics: https://www.med.unc.edu/pgc/download-results/
- HapMap3 SNP-list: https://data.broadinstitute.org/alkesgroup/LDSCORE/w_hm3.snplist.bz2
- GTEx Portal: https://gtexportal.org/home/datasets
- Software:
  - FUSION repository: https://github.com/gusevlab/fusion_twas
  - FeaturePred repository: https://github.com/opain/Predicting-TWAS-features
  - eQTL_to_TWAS repository: https://github.com/opain/eQTL_to_TWAS

## Acknowledgements

We thank Jian Zeng and Michael J. Gandal for providing helpful comments on the manuscript prior to submission. We also thank the eQTLGen consortium for making their results publicly available and providing us with results independent of GTEx for our study.

The authors acknowledge use of the research computing facility at King’s College London, Rosalind (https://rosalind.kcl.ac.uk), which is delivered in partnership with the NIHR Maudsley BRC, and part-funded by capital equipment grants from the Maudsley Charity (award 980) and Guy’s & St. Thomas’ Charity (TR130505). The views expressed are those of the authors and not necessarily those of the NHS, the NIHR or the Department of Health and Social Care.

The Genotype-Tissue Expression (GTEx) Project was supported by the Common Fund of the Office of the Director of the National Institutes of Health (commonfund.nih.gov/GTEx). Additional funds were provided by the NCI, NHGRI, NHLBI, NIDA, NIMH, and NINDS. Donors were enrolled at Biospecimen Source Sites funded by NCI\Leidos Biomedical Research, Inc. subcontracts to the National Disease Research Interchange (10XS170), Roswell Park Cancer Institute (10XS171), and Science Care, Inc. (X10S172). The Laboratory, Data Analysis, and Coordinating Center (LDACC) was funded through a contract (HHSN268201000029C) to The Broad Institute, Inc. Biorepository operations were funded through a Leidos Biomedical Research, Inc. subcontract to Van Andel Research Institute (10ST1035). Additional data repository and project management were provided by Leidos Biomedical Research, Inc.(HHSN261200800001E). The Brain Bank was supported supplements to University of Miami grant DA006227. Statistical Methods development grants were made to the University of Geneva (MH090941 & MH101814), the University of Chicago (MH090951,MH090937, MH101825, & MH101820), the University of North Carolina – Chapel Hill (MH090936), North Carolina State University (MH101819),Harvard University (MH090948), Stanford University (MH101782), Washington University (MH101810), and to the University of Pennsylvania (MH101822). The datasets used for the analyses described in this manuscript were obtained from dbGaP at http://www.ncbi.nlm.nih.gov/gap through dbGaP accession number phs000424.v8.p2.

## Supplementary Information

### Pseudovalidation of TWAS models

In the setting of using eQTL summary statistics, formal validation of each model is not possible, so we tested a TWAS-based pseudovalidation approach to infer which model best predicted expression of a given gene.

On average, when using pseudovalidation to select the best model, the predicted-observed correlation was 0.026 lower than the best model identified by testing in GTEx (Figure S5). Between pseudovalidation and external validation in GTEx, the average rank correlation of methods was 0.097, and pseudovalidation selected the best model for 16% of genes. Confusion matrices were created to determine predictive utility of pseudovalidation for each model (Figure S5). There was a substantial difference in performance of pseudovalidation across models, with a very low sensitivity for the top1 model despite often performing best. The bias towards more complex models may be a result of overfitting due to the use of the eQTL summary statistics to both train the TWAS models and test their predictive performance. The median correlation between predicted and observed expression for models selected by pseudovalidation was 0.024, indicating using pseudovalidation to select the model for each gene will lead to poorer prediction of expression than any one method applied to all genes.

We also evaluated pseudovalidation results for FUSION-YFS based models, comparing the reported 5-fold cross-validation variance explained in the YFS sample. The concordance between pseudovalidation and formal validation was similar in YFS as it was in eQTLGen (Figure S6).

### SNP-based heritability of gene expression

We compared the SNP-based heritability of the 561 genes on chromosome 22 based on GTEx v8 whole blood eQTL summary statistics to the SNP-based heritability reported by FUSION using GREML and the individual-level GTEx v8 data (Figure S7, Table S3). We also highlight the number of genes for which each method converged when estimating SNP-based heritability. The summary statistic method providing SNP-based heritability estimates most similar to GREML was SBayesR, with a correlation of 0.902 between SBayesR and GREML SNP-based heritability estimates. SBayesR successfully converged for 560 out of 561 genes. SNP-based heritability estimates from SBayesR-robust were highly correlated with those of SBayesR (0.975), but the correlation with GREML estimates was lower (0.863) and the method only converged for 285 genes. SNP-based heritability estimates from LDSC were highly inaccurate, with most estimates outside the 0−1 range, and the correlation with GREML estimates was 0.556. The poor performance of LDSC was expected as it is not designed to accurately estimates SNP-based heritability within a specific locus.

FUSION traditionally only generates TWAS models for genes with a SNP-based heritability *p*<0.01. Of the 561 genes considered, GREML identified 301 genes with significant SNP-based heritability, which was more SNP-based heritable genes than any summary statistic-based method estimating SNP-based heritability. Of the summary statistic-based methods, SBayesR identified the largest number of genes with a statistically significant SNP-based heritability (N=275) and identified the most SNP-based heritable genes that were also identified as SNP-based heritable by GREML (N=271).

Concordant with the correlation analysis with GREML estimates, the other methods were also less congruent with GREML in terms of overlapping SNP-based heritable genes.

We also evaluated an alternative approach for defining inclusion criteria for genes in the TWAS, including genes if they had a genome-wide significant eQTL present, or mBAT-combo estimated a significant gene association statistic (FDR < 0.05). Using these criteria identified the largest of number of genes (N=302) compared to other summary statistic-based methods, and the largest number genes identified as significant by GREML (N=287). This method converges for all genes.

## Supplementary Figures

**Figure S1.**
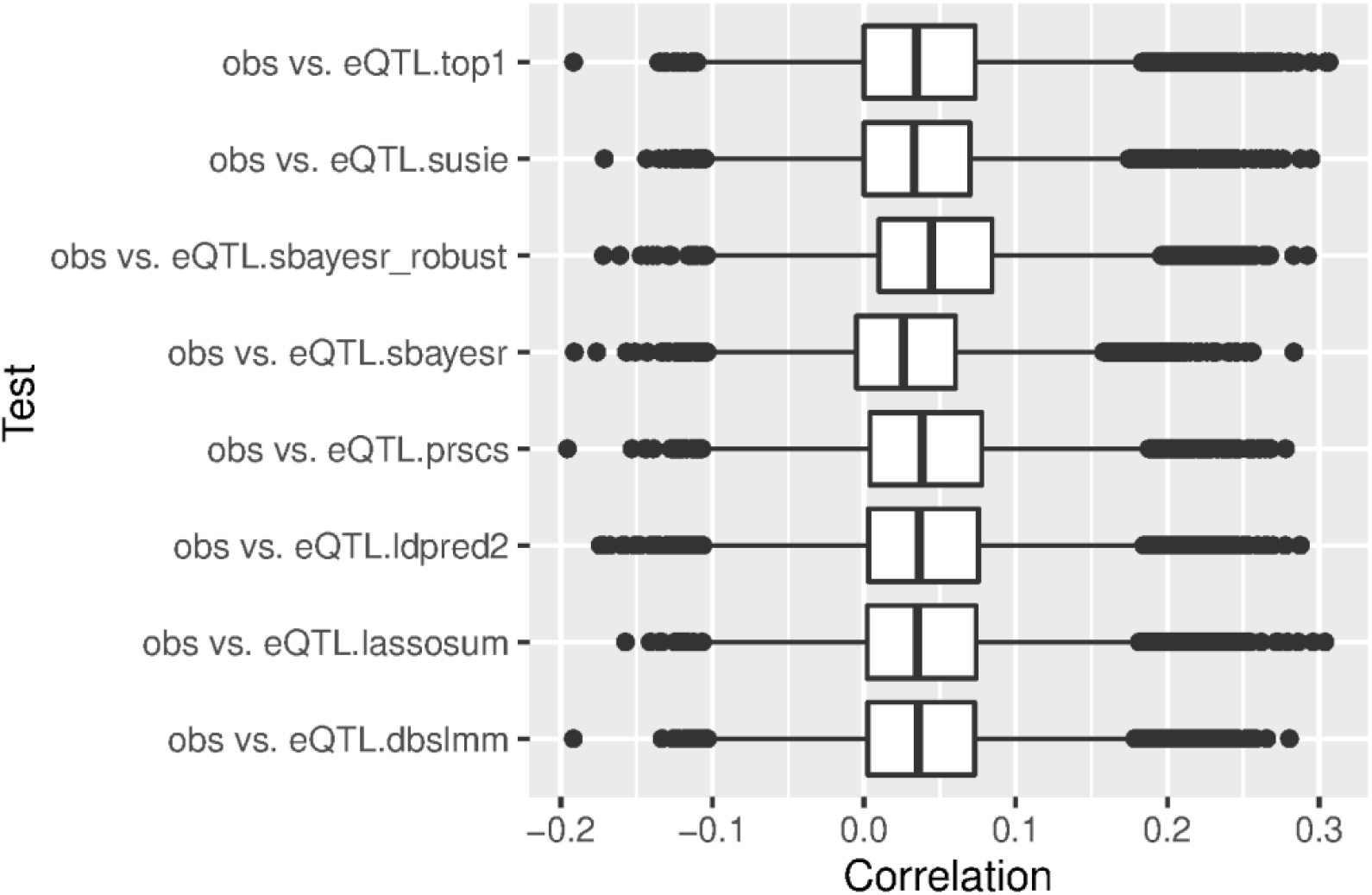
Median predicted-observed correlation using eQTLGen summary statistics and GTEx v8 target sample.

**Figure S2.**
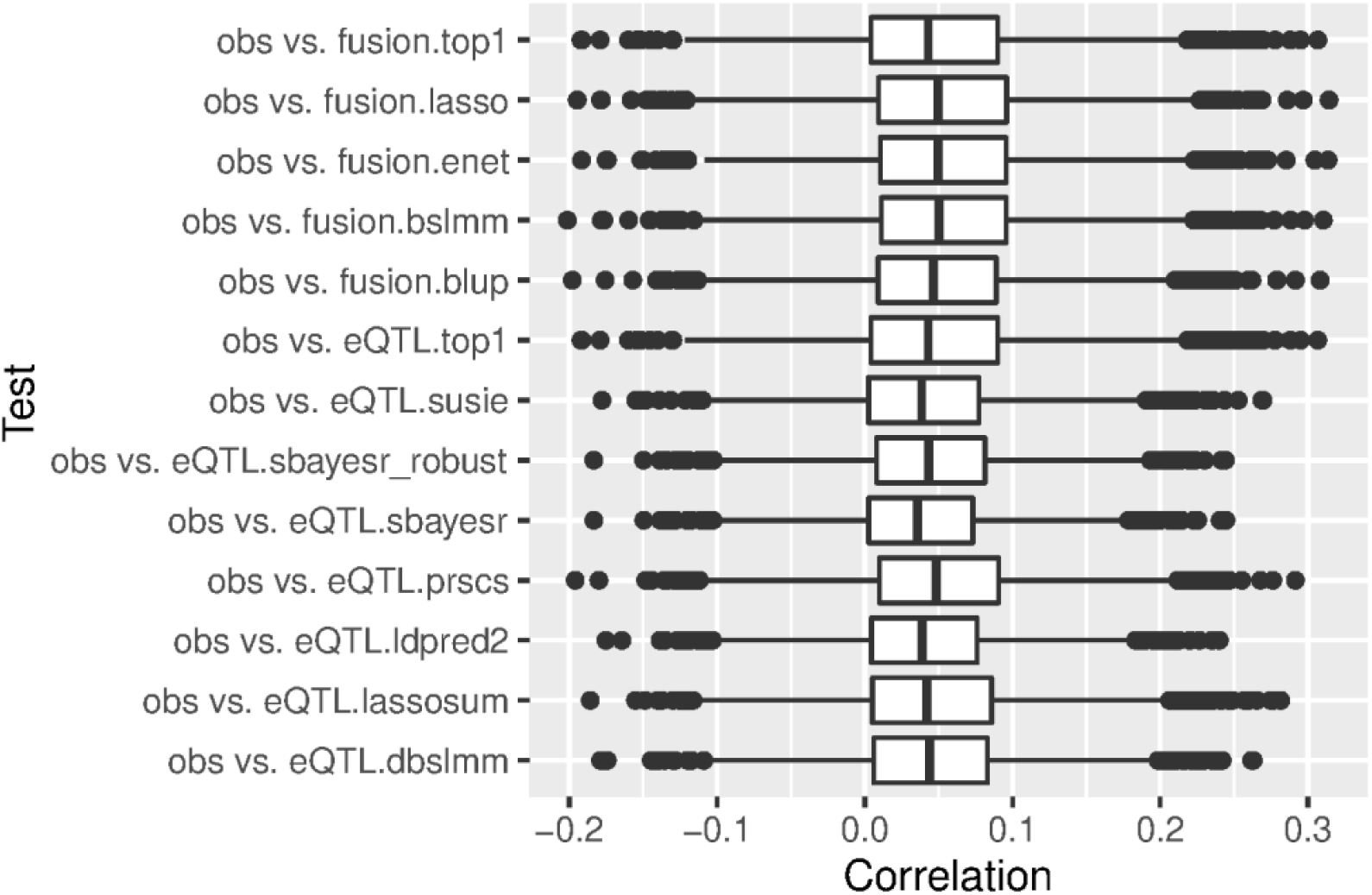
Median predicted-observed correlation using YFS summary statistics and GTEx v8 target sample.

**Figure S3.**
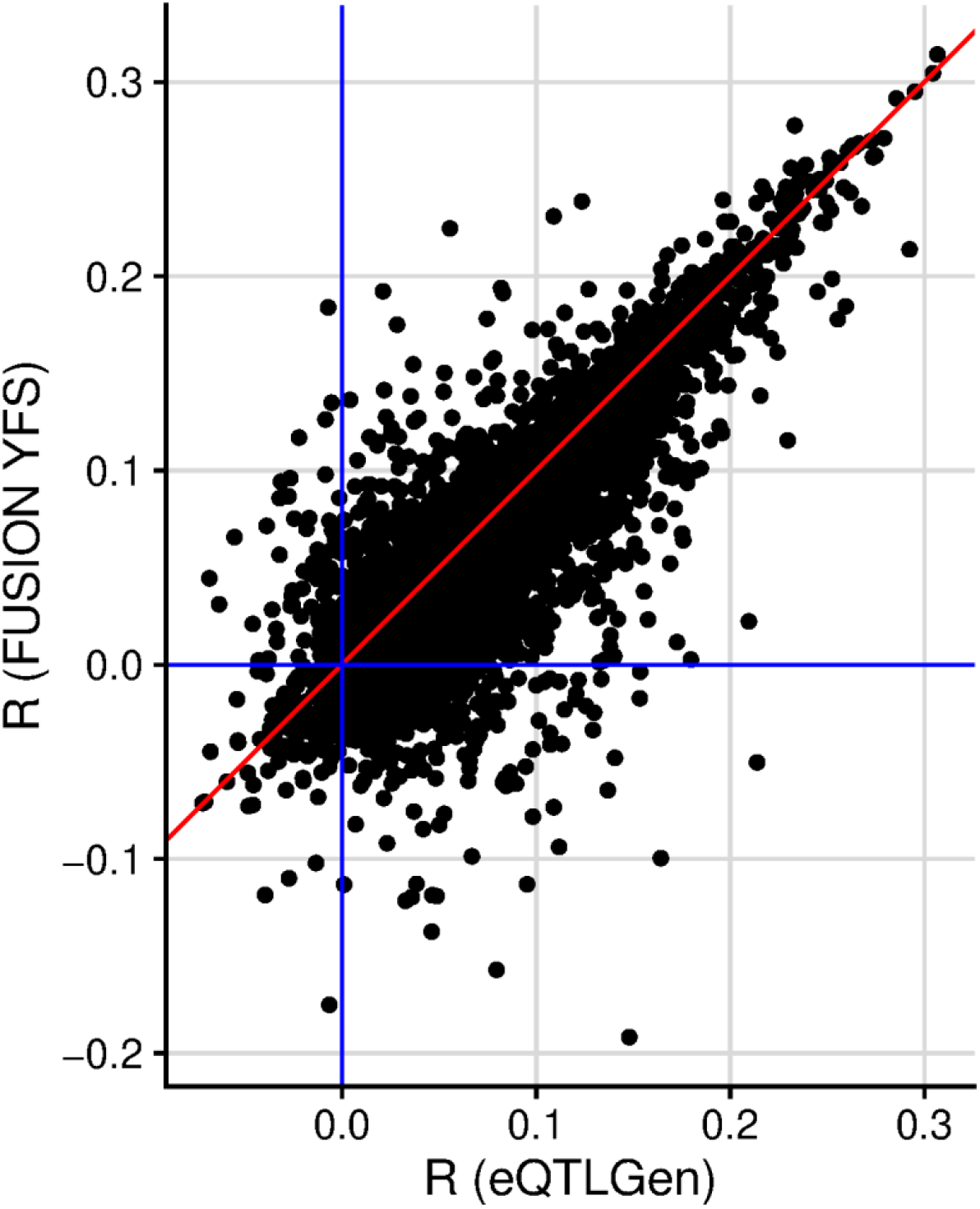
Predicted-Observed correlation for the best model within FUSION YFS and eQTLGen datasets.

**Figure S4.**
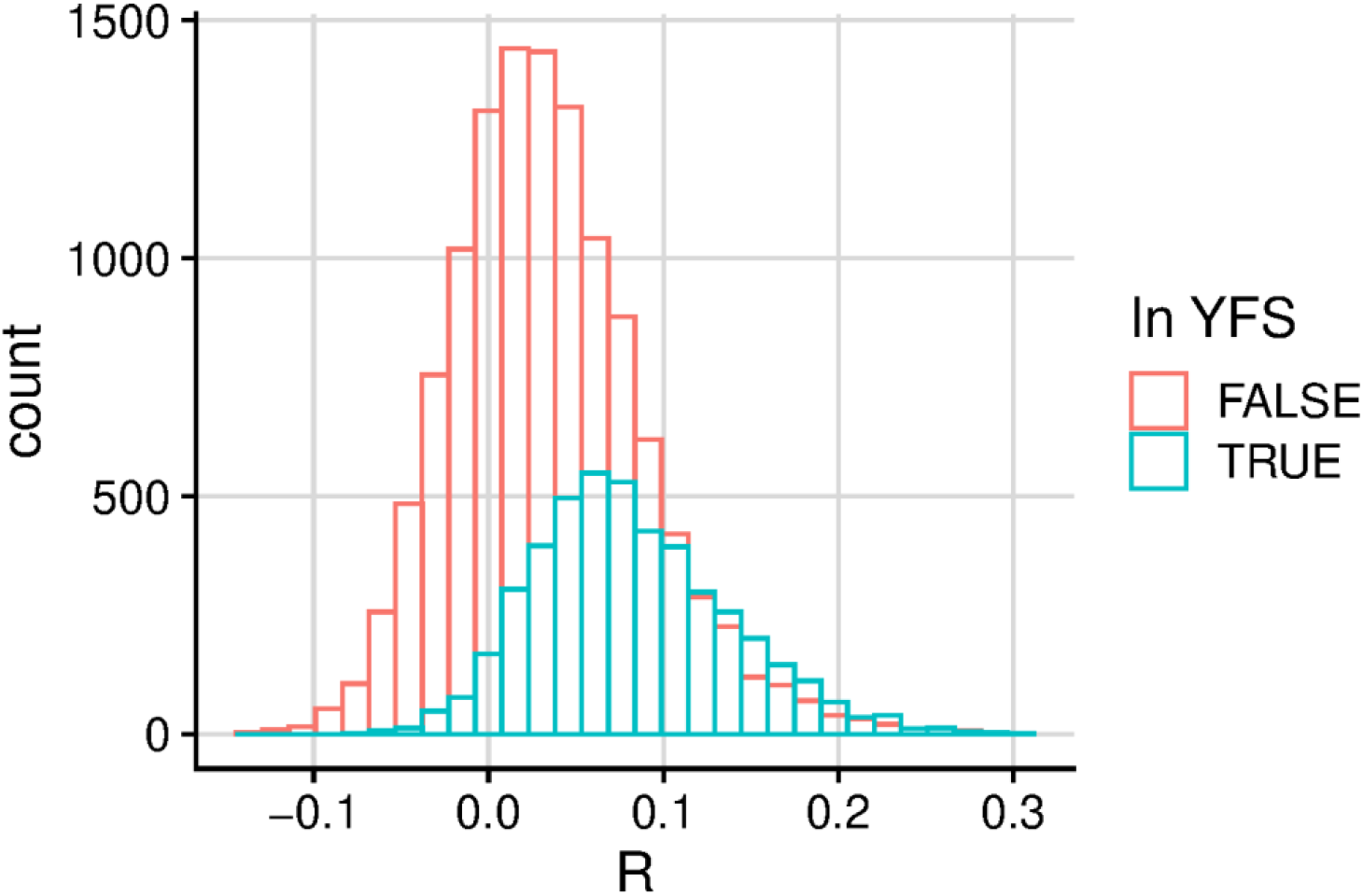
Distribution of predicted-observed correlation of genes in eQTLGen stratified by their presence in the FUSION YFS panel.

**Figure S5.**
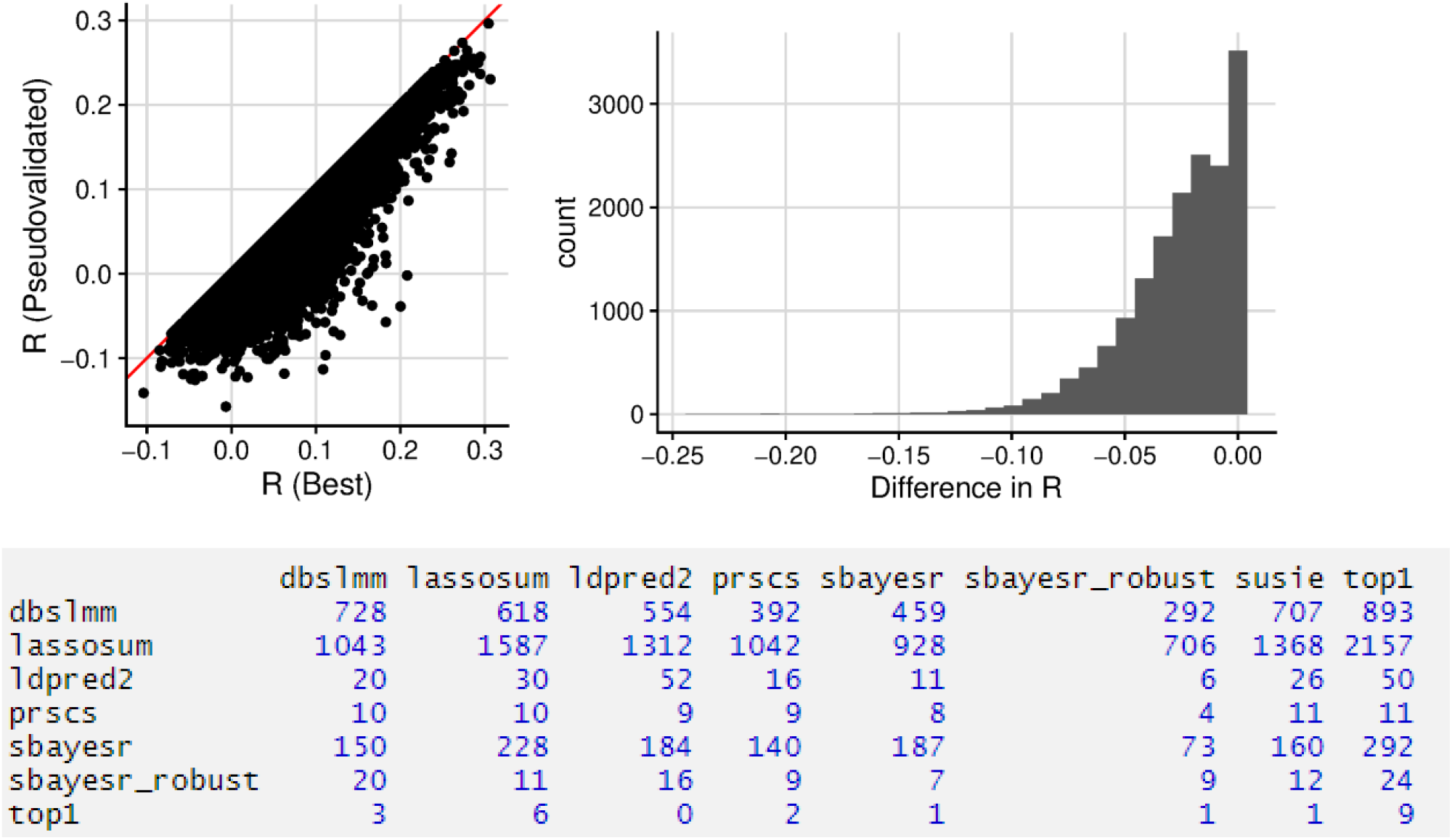
Comparison of predicted-observed correlation in the GTEx v8 sample of the best model identified in GTEx v8 and the best model identified using pseudovalidation. Confusion matrix shows true best model on x-axis, and predicted best model on y-axis.

**Figure S6.**
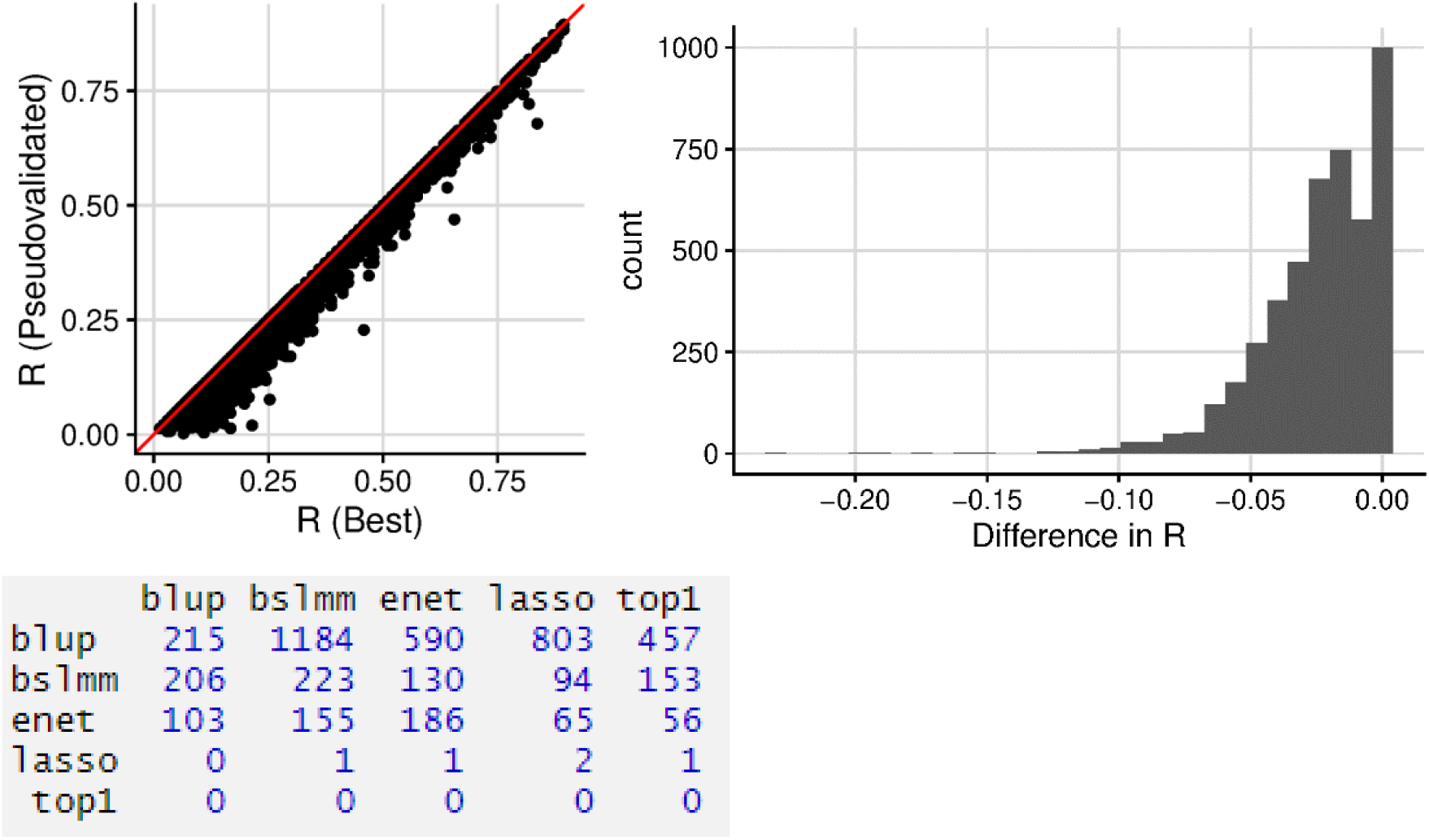
Comparison of predicted-observed correlation in the YFS sample of the best model identified using 5-fold cross validation in the YFS sample and the best model identified using pseudovalidation. Confusion matrix shows true best model on x-axis, and predicted best model on y-axis.

**Figure S7.**
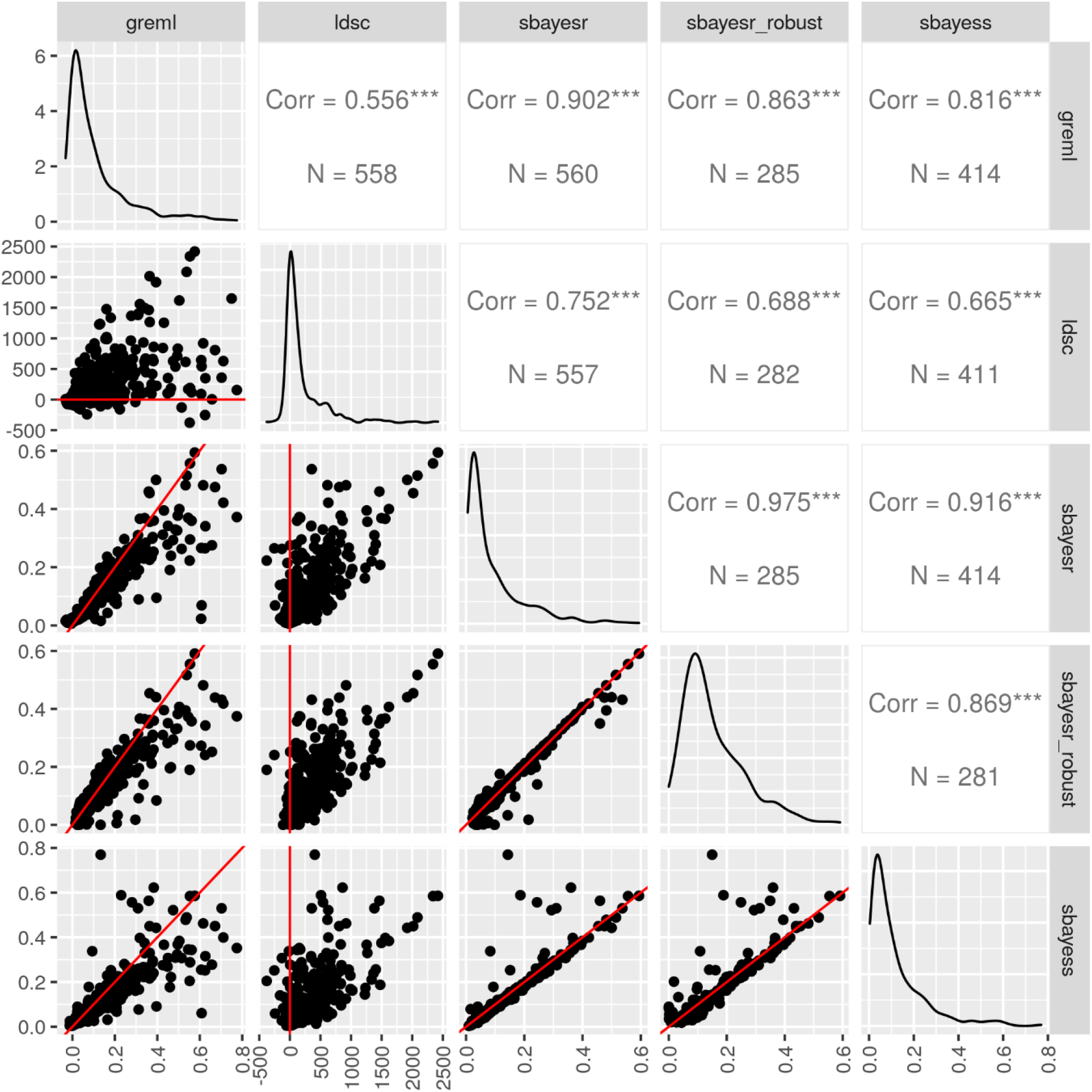
Pairwise comparison of SNP-based heritability estimates. Lower triangle shows scatterplot of SNP-based heritability estimates from each method with the red line indicating a correlation of 1. The diagonal elements are density plots showing the distribution of SNP-based heritability estimates for each method. The upper triangle shows the Pearson correlation between SNP-based heritability estimates and the number of estimates available for both methods.

## Supplementary Tables

**Table S1.** Predicted-observed correlation using FUSION-YFS and eQTL-YFS TWAS models, and GTEx v8 target sample. N valid and % valid indicate the number and percentage of genes that had a predicted-observed correlation > 0.1. Columns labelled ‘eQTL’ only compare the performance of models derived using eQTL summary statistics. N sig. top (eQTL) indicates the number of genes for which an eQTL summary statistic-based method showed a statistically significant improvement in prediction over all other eQTL summary statistic-based methods.

| Method | N valid | Median r | % valid | N top | Median r top | % top | N top (eQTL) | Median r top (eQTL) | % top (eQTL) | N sig. top (eQTL) |
| --- | --- | --- | --- | --- | --- | --- | --- | --- | --- | --- |
| BSLMM (FUSION) | 1065 | 0.05 | 23.00% | 394 | 0.093 | 8.50% | NA | NA | NA | NA |
| Elastic Net (FUSION) | 1067 | 0.05 | 23.00% | 397 | 0.09 | 8.60% | NA | NA | NA | NA |
| Lasso (FUSION) | 1071 | 0.049 | 23.10% | 406 | 0.088 | 8.80% | NA | NA | NA | NA |
| PRS-CS (eQTL) | 963 | 0.048 | 20.80% | 244 | 0.084 | 5.30% | 483 | 0.084 | 10.40% | 7 |
| BLUP (FUSION) | 952 | 0.047 | 20.50% | 412 | 0.074 | 8.90% | NA | NA | NA | NA |
| DBSLMM (eQTL) | 834 | 0.044 | 18.00% | 346 | 0.066 | 7.50% | 511 | 0.065 | 11.00% | 5 |
| SBayesR-robust (eQTL) | 628 | 0.043 | 13.60% | 107 | 0.067 | 2.30% | 174 | 0.068 | 3.80% | 6 |
| top1 (FUSION) | 959 | 0.043 | 20.70% | 3 | 0.003 | 0.10% | NA | NA | NA | NA |
| Top1 (eQTL) | 958 | 0.043 | 20.70% | 707 | 0.072 | 15.30% | 1169 | 0.076 | 25.30% | 77 |
| lassosum (eQTL) | 861 | 0.042 | 18.60% | 430 | 0.061 | 9.30% | 754 | 0.069 | 16.30% | 26 |
| LDpred2 (eQTL) | 657 | 0.039 | 14.20% | 362 | 0.053 | 7.80% | 482 | 0.052 | 10.40% | 23 |
| SuSiE (eQTL) | 707 | 0.039 | 15.30% | 599 | 0.062 | 12.90% | 759 | 0.064 | 16.40% | 29 |
| SBayesR (eQTL) | 534 | 0.035 | 11.50% | 222 | 0.052 | 4.80% | 297 | 0.052 | 6.40% | 10 |

**Table S2.** Summary of results for FUSION-YFS and eQTL-YFS based TWAS of schizophrenia, using different model selection criteria and aggregation methods. N test = Number of genes with non-missing TWAS Z scores in the output; N sig. = Number of significant genes after Bonferroni correction; N sig. overlap with FUSION (CV.R2) = Number of significant genes also significant in the FUSION-YFS (CV.R2) TWAS; CV.R2 = the model was selected with the highest variance explained in YFS (default behaviour in FUSION); PseudoVal = the model was selected with the highest Pseudovalidation Z score; TWAS.P = the model was selected with the smallest TWAS p-value (most significant); FUSION.O = results for each model was aggregated using the FUSION OMNIBUS test; ACAT.O = results for each model was aggregated using the ACAT-O test; SNP-h^2^ = only genes with a statistically significant SBayesR SNP-based heritability were retained; mBAT-combo, only genes with either a significant mBAT-combo gene association or genome-wide significant eQTL were retained.

| Approach | N test | N sig. | N sig. coloc. | N sig. overlap with FUSION (CV.R2) | N sig. coloc. overlap with FUSION (CV.R2) |
| --- | --- | --- | --- | --- | --- |
| FUSION (CV.R2) | 4685 | 93 | 21 | 93 | 21 |
| eQTL (PseudoVal) | 4685 | 75 | 12 | 58 | 12 |
| eQTL (SNP-h<sup>2</sup>; PseudoVal) | 4028 | 49 | 9 | 39 | 9 |
| eQTL (mBAT-combo; PseudoVal) | 4677 | 75 | 12 | 58 | 12 |
| eQTL (ACAT.O) | 4685 | 162 | 28 | 87 | 18 |
| eQTL (SNP-h<sup>2</sup>; ACAT.O) | 4028 | 104 | 22 | 64 | 15 |
| eQTL (mBAT-combo; ACAT.O) | 4677 | 162 | 28 | 87 | 18 |
| eQTL (FUSION.O) | 4683 | 605 | 33 | 82 | 17 |
| eQTL (SNP-h<sup>2</sup>; FUSION.O) | 4026 | 487 | 27 | 60 | 14 |
| eQTL (mBAT-combo; FUSION.O) | 4675 | 604 | 33 | 82 | 17 |

**Table S3.** Analysis of 561 genes on chromosome 22. Compares each method in terms of the number of genes for which each SNP-based heritability method converged successfully (N converge), the number of genes identified with statistically significant SNP-based heritability (N significant), and the number of SNP-based heritable genes identified in common with GREML. The number of genes identified as significant by mBAT-combo or containing a genome-wide wide significant variant are also included. Note. mBAT-combo refers to selecting genes with a mBAT-combo FDR < 0.05 or at least one genome-wide significant eQTL.

| Method | N converge | N significant | N GREML overlap |
| --- | --- | --- | --- |
| GREML | 561 | 301 | 301 |
| LDSC | 558 | 256 | 184 |
| SBayesR | 560 | 275 | 271 |
| SBayesR-robust | 285 | 263 | 243 |
| SbayesS | 414 | 247 | 244 |
| mBAT-combo | 561 | 302 | 287 |

**Table S4.**
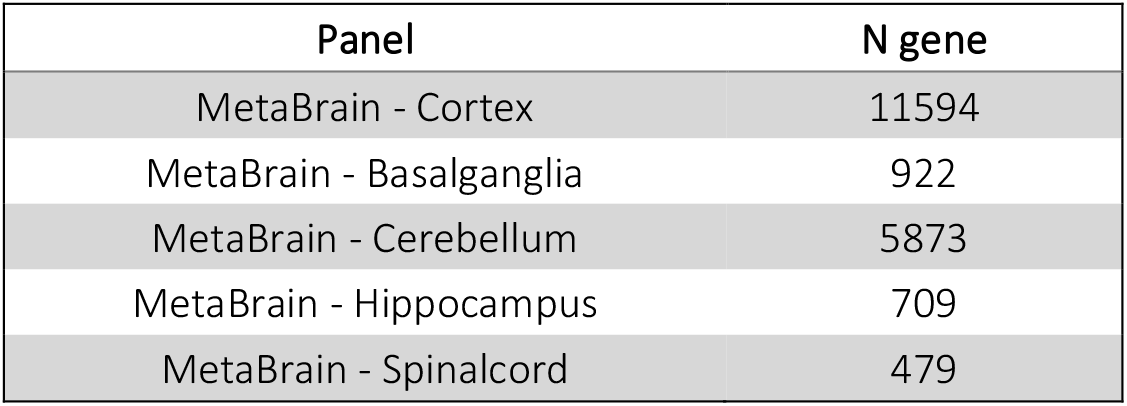
Number of genes for each MetaBrain tissue with either an FDR significant mBAT-combo association or a genome-wide significant eQTL.

